# Reducing RF-induced Heating near Implanted Leads through High-Dielectric Capacitive Bleeding of Current (CBLOC)

**DOI:** 10.1101/456533

**Authors:** Laleh Golestanirad, Leonardo M Angelone, John Kirsch, Sean Downs, Boris Keil, Giorgio Bonmassar, Lawrence L Wald

## Abstract

Patients with implanted medical devices such as deep brain stimulation or spinal cord stimulation are often unable to receive magnetic resonance imaging (MRI). This is because once the device is within the radiofrequency (RF) field of the MRI scanner, electrically conductive leads act as antenna, amplifying the RF energy deposition in the tissue and causing possible excessive tissue heating. Here we propose a novel concept in lead design in which 40cm lead wires are coated with a ~1.2mm layer of high dielectric constant material (155 < *ε_r_* < 250) embedded in a weakly conductive insulation (*σ* = 20 *S/m*). The technique called High-Dielectric Capacitive Bleeding of Current, or CBLOC, works by forming a distributed capacitance along the lengths of the lead, efficiently dissipating RF energy before it reaches the exposed tip. Measurements during RF exposure at 64 MHz and 123 MHz demonstrated that CBLOC leads generated 20-fold less heating at 1.5 T, and 40-fold less heating at 3 T compared to control leads. Numerical simulations of RF exposure at 297 MHz (7T) predicted a 15-fold reduction in specific absorption rate (SAR) of RF energy around the tip of CBLOC leads compared to control leads.

## I. INTRODUCTION

There is a steady growth in the use of implantable electronic devices for therapeutic applications in the US and globally. As of 2012, there were more than 3 million patients with permanent pacemakers, cardioverter defibrillators, or cardiac resynchronization therapy devices in the US [1], with the number growing by 80,000 annually [2]. In the field of neuromodulation, the market for spinal cord stimulation (SCS), sacral nerve stimulation, and deep brain stimulation (DBS) is expected to grow at an annual rate greater than 17% from 2016 to 2023 [3]. Factors such as aging population, increasing prevalence of cardiovascular and neurological diseases, expanded target applications, and new indications of use are among those driving such steep growth.

For a majority of neurologic, cardiac and musculoskeletal disorders, magnetic resonance imaging (MRI) is the diagnostic modality of choice because of its excellent soft tissue contrast and non-invasive nature. It is estimated that 50-75% of patients with cardiovascular implantable devices may need to undergo MRI over their lifetime for non-cardiac or cardiac indications [4], with many patients requiring repeated examinations [5]. Similarly, patients with neuromodulation devices such as DBS greatly benefit from MRI exams, both for target verification and for post-operative monitoring of treatment-induced changes in the function of affected brain networks [6]. Unfortunately, however, the interaction of radiofrequency (RF) fields of MRI transmitters with implanted leads results in safety hazards that severely limit the post-operative accessibility of MRI for patients with implanted conductive leads.

One major safety concern is RF-induced heating of tissue due to the “antenna effect” of the leads, where the electric field of the MRI transmit coil couples with the elongated conductive leads and amplifies the specific absorption rate (SAR) of the RF energy in the tissue [7, 8]. Such SAR amplification can cause excessive tissue heating and potential tissue damage as underscored by several injuries reported worldwide [9, 10]. As a result, the conditions under which patients with implanted leads can receive an MRI are restrictive. For DBS patients for example, only field strength of 1.5 T is permitted (excluding centers that only have a higher field strength); only pulse sequences with a whole-head SAR of 0.1 W/kg (approximately 30-fold below the FDA-approved clinical level) or effective B1rms<2μT are allowed, and the current state-of-the-art MRI parallel transmit coils are contraindicated [11]. Similar restrictions apply to MR Conditional SCS systems and neuromodulation systems for chronic pain [12].

Efforts to alleviate the problem of implant-induced tissue heating during MRI can be classified into three main categories: those that aim to modify the imaging hardware to make it less interactive with conductive implants, those that modify the implant structure and material to reduce the antenna effect, and those that – through surgical planning - modify the implant trajectory to reduce the coupling and the antenna effect. Hardware modification is a relatively new concept and its scientific literature mostly consists of proof-of-concept studies and early-stage prototypes. Examples include the use of dual-drive birdcage coils to generate steerable low-E field regions that coincide with the implant [13, 14], the introduction of rotating linear birdcage coils that allow individual patient adjustments for low SAR imaging [15, 16], and parallel transmit systems that produce implant-friendly modes [17, 18].

Alteration of the lead geometry on the other hand has a much older history as attested by the plethora of patents published in the past fifteen years [19–26]. This body of work mostly consists of techniques that aim to increase the lead’s impedance to reduce induced RF currents. More recent contributions suggested the use of resistive tapered stripline to scatter the RF energy along the length of the lead and reduce its concentration at the tip [27], use of reverse or overlapping windings [28, 29], use of external traps which couple to lead wires and take the RF energy away from internal wires [30], and the use of conductive pins to connect lead wires to the tissue and shunt induced currents [31]. Finally, the role of surgical planning to optimize the routing of implanted leads to reduce the heating at the tip has been explored [32, 33]. Despite these efforts however, the number of MR Safe or MR Conditional implantable leads remains limited and the need to develop effective strategies to safeguard patients is more urgent than ever.

This paper introduces a novel lead design to improve RF safety of implantable electronic devices in MRI environment. The technique termed High-Dielectric Capacitive Bleeding of Current, or CBLOC is based on a distributed capacitive dispersion of RF energy along the length of the lead which significantly reduces the energy concentration at the exposed tip of the implant. This is achieved by using a thin layer (~1.2mm) of high dielectric constant (HDC) material to coat the lead wires, and then providing a conductive path to the tissue by embedding them within partially conductive Carbone-doped silicon tubing (*σ* = 20 *S*/*m*). This allows a distributed capacitance to be formed between the wire, HDC material, and the conductive tubing. This capacitive element will then shunt the RF energy along the length of the lead in the form of displacement currents and thus, reduces the RF heating at the tip. We show that this technique is remarkably effective, reducing the temperature rise at the tip of the lead by up to 20-fold. Furthermore, unlike techniques based on resonant RF traps and filters, the CBLOC performance is independent of the RF frequency, showing a substantial heat reduction effect across 1.5 T, 3 T and 7 T systems.

This manuscript is laid out as follows: Section II gives a theoretical background on the phenomenology of RF heating of implants in MRI environment. The fundamentals of methodologies to reduce RF heating of implanted leads during MRI are reviewed, and the proposed CBLOC technique is described. Section III gives the details of experimental setup, lead construction, and temperature measurements during MR RF exposure at 64 MHz (1.5 T) and 123 MHz (3T). Finally, Section IV presents results of numerical modeling confirming the same trend in the reduction of RF heating as observed in experimental measurements at 1.5 T, 3 T and predicting the behavior of CBLOC leads at 7 T.

## II. THEORY

The phenomenology of RF heating in the presence of linear conductive structures in the MRI environment has been studied in a number of theoretical contributions [8, 34–37]. The consensus is that the tangential component of the electric field, ***E***_tan_, along the length of the wire acts as a local voltage source generating RF currents on the lead. For an insulated wire with an exposed tip, scattered fields will arise at the tip to enforce the boundary condition ***E***_tan_=0. These scattered fields generate electric currents that produce heating when dissipated in the tissue. Figure 1A gives the general circuit model of an insulated lead wire inside an MRI scanner, assuming the lead is a good conductor with low resistivity (R_Lead_≈0). From the model, one obvious approach to reduce RF currents is to increase the lead’s impedance at the desired frequency. This technique has been the subject of numerous designs, including the use of tightly wound loops to increase the inductance of the wire [29, 38] and insertion of RF traps [39] or bands-top filters [40] to introduce a high impedance component along the length of the lead.

**Figure 1:**
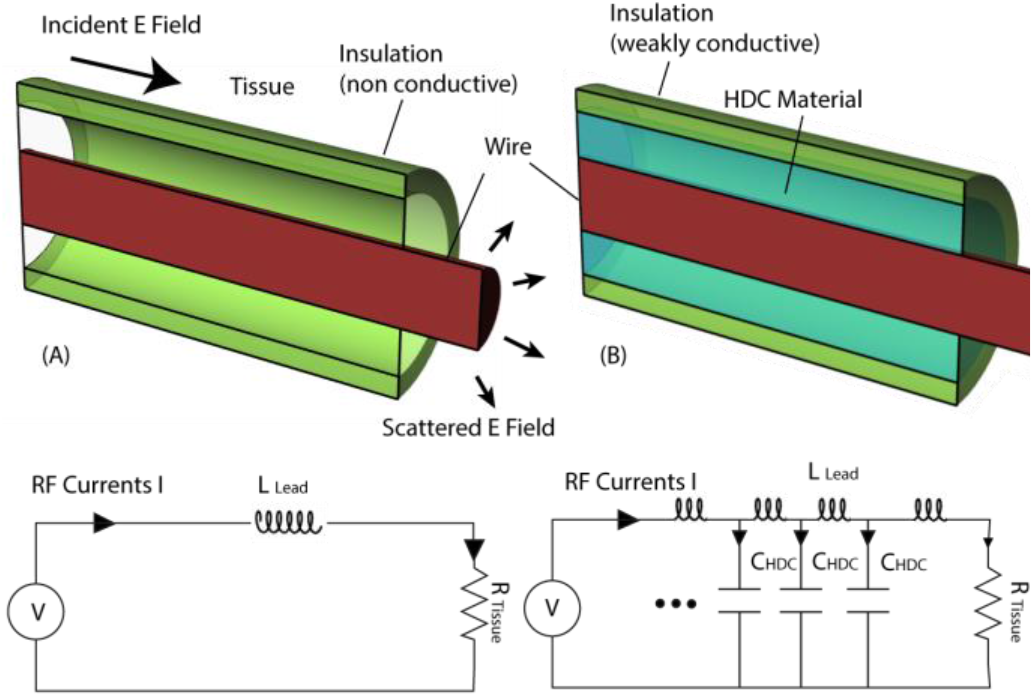
(A) Equivalent circuit of a lead in MRI environment. The tangential component of the electric field acts as a voltage source inducing currents on the lead (modelled as an indicator) which will then be dissipated in the lossy tissue. (B) The equivalent circuit of a lead with an HDC core contained in weakly conductive insulation.

Another possible approach to reduce the magnitude of RF currents at the tip is to shunt the currents along the length of the lead through capacitive elements. A suggested implementation of this approach is, for example, the insertion of intermittent pins connecting internal wires to the conductive tissue [31]. Such techniques, however, have not been widely explored due to difficulties in design and implementation of lumped capacitive components. Our proposed method is to implement a distributed capacitance along the length of the lead through a thin layer of HDC material embedded in a weakly conductive insulator. Figure 1B gives the equivalent circuit model of such a lead. Theoretically, this structure can effectively shunt the RF energy along the length of the lead while leaving DC currents associated with the therapeutic function of the device unperturbed.

To ensure that the method is effective, the HDC material should be either in direct contact with the conductive tissue (if the material is solid) or in contact with a conductive tubing (as in our case where the dielectric material is in the form of a paste which needs to be contained.) The reason for this can be better appreciated considering the theory of partially-filled capacitors. Figure 2A gives the equivalent circuit of a lead with HDC-coated wire embedded within a conductive insulation (e.g., medical grade carbon-doped silicone). The distributed capacitor C1 is formed between the lead wire (first conductor), the HDC layer, and the tubing (second conductor). In contrast, Figure 2B shows the HDC-coated wire encapsulated within a nonconductive tubing such as polyurethane commonly used as insulation in medical implants. In this case, the total capacitance C is the series combination of the two partially filled cylindrical capacitors as 1/C=1/C_1_+1/C_2_ [41]. Because the outer insulator has a much lower dielectric constant (*ε_r_* ≅ 3) the effect of the HDC coating will be negated in this case. For this reason, it is critical to use the HDC coating in contact with a weakly conductive insulation to provide a path for the RF energy to dissipate.

**Figure 2:**
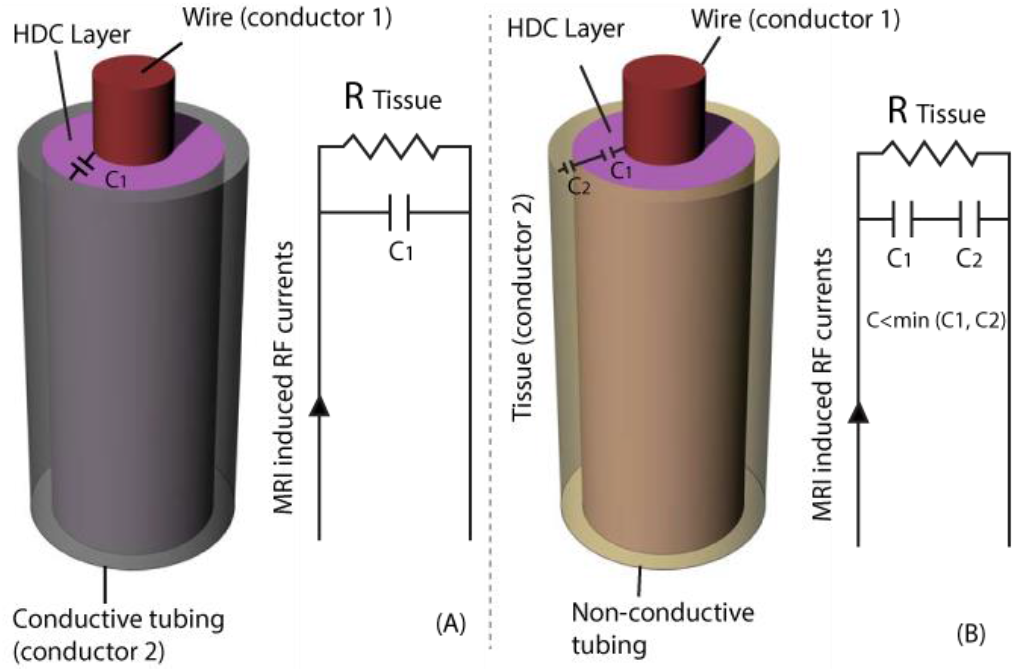
Schematic and equivalent circuit of a lead with an HDC core (A) contained in conductive tubing and (B) contained in non-conductive rubber tubing.

## III. EXPERIMENTS

To test out the theory outlined in Section II, experiments were conducted by measuring MRI-induced temperature rise at the tip of different lead constructs at 1.5 T and 3 T scanners. This section outlines the details of experimental setup, lead construction, and measurement results.

### A. Material and Construction of the Leads

#### 1) HDC paste

A high dielectric paste was made by mixing Barium titanate (BaTiO3) powder (Sigma-Aldrich, St Louis, MO) with distilled, de-ionized water until a saturated suspension was obtained as described in [42]. Sodium polyacrylate (Sigma-Aldrich, St Louis, MO) was used as a chemical dispersant, added one drop at a time to the paste until it became unsaturated and could absorb more dry powder. The dielectric constant of the paste was then measured using a dielectric probe kit (85070E, Agilent Technologies, Santa Clara, CA) and a network analyzer. Figure 3 gives the relative permittivity of the paste as measured by the network analyzer, showing *ε_r_* = 250 at 64 MHZ (1.5 T), *ε_r_* = 204 at 123 MHz (3 T), and *ε_r_* = 155 at 279 MHz (7 T).

**Figure 3:**
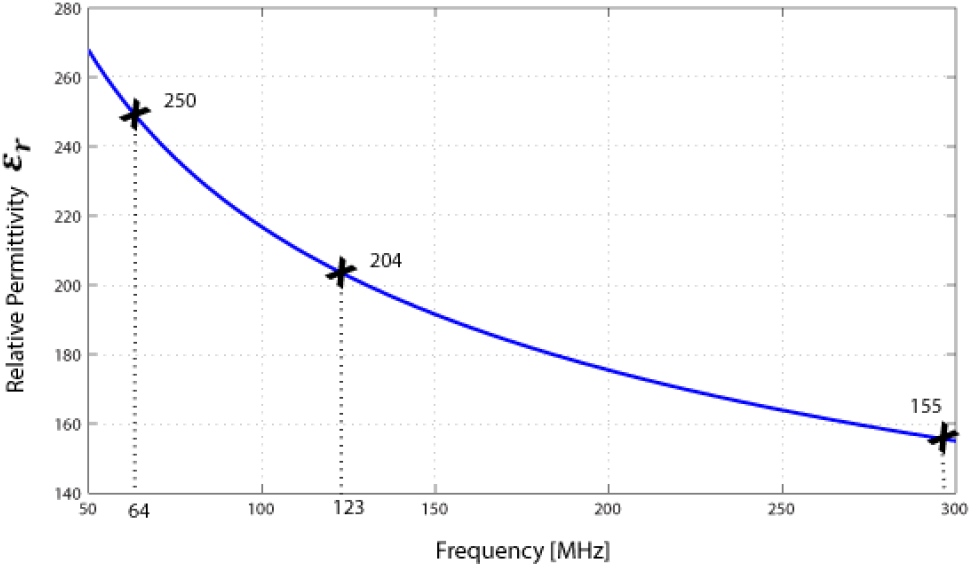
Relative permittivity of the HDC paste measured at 64 MHz (1.5 T) and 123 MHz (3 T), and 297 MHz (7 T) using the dielectric probe kit and a network analyzer.

#### 2) Lead wires and tubing

Leads were constructed from 40 cm Ga22 varnished copper wires (magnet wire 7588k79, 0.0021” varnish thickness McMaster-Carr, Elmhurst, IL), with 1 cm of the insulation removed at one end to expose the tip. Wires were inserted into either a rubber tubing (*ε_r_* = 3, inside diameter (ID) = 3 mm, outside diameter (OD) = 4 mm) or a conductive silicone tubing (*σ* = 20 *S/m*, *ε_r_* = 3.5, ID=2.7mm, OD=4mm). Wires were stretched between the two ends of the tubing and passed through rubber stoppers to seal and support (Figure 4).

**Figure 4:**
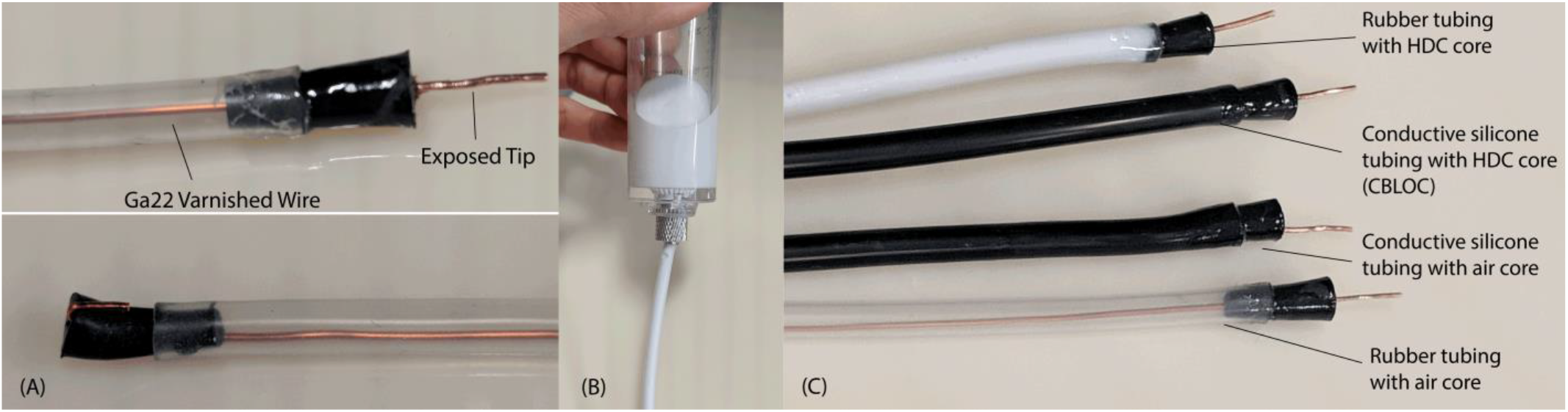
(A) Ga22 varnished wires were inserted into tubing and passed through rubber stoppers at both ends to seal and support. (B) The HDC paste being injected into tubing using a Ga14 syringe. (C) Different lead configurations used in the experiments.

To examine the theory laid out in Section II, we built four types of lead with (*a*) wires inserted into an empty rubber tubing (control lead), (*b*) wires inserted into a rubber tubing filled with the HDC paste, (*c*) wires inserted into an empty conductive silicone tubing, and (*d*) wires inserted into a silicone conductive tubing filled with the HDC paste (CBLOC lead). We hypothesized that the CBLOC lead (lead (*d*)) significantly reduces the temperature rise in the tissue compared to the control lead (lead (*a*)), whereas configurations (*b*) and (*c*) will have little effect on the heating. Note that lead (*b*) has the equivalent circuit shown in Figure 2B, in which the effective distributed capacitance is the series combination of capacitors formed between the HDC layer (C_1_) and the plastic insulation (C_2_) with the value being smaller than both C_1_ and C_2_.

### B. RF Exposure and Temperature Measurements

An anthropomorphic head phantom was designed and 3D-printed based on the structural MRI of a healthy volunteer (Figure 5). The mold was composed of two sagittal parts connected through a rim along the central sagittal plane. Phantom dimensions were approximately 16cm ear-to-ear and 27cm from top of head to bottom of neck. The phantom was filled with agarose-doped saline solution that mimicked the electrical and thermal properties of biological tissues (*ε_r_* ≅ 70, *σ* ≅ 1 *S*/*m*, *C_p_* = 4150 J/kg°C). Gel recipe and construction method is given elsewhere [15]. A relatively high percentage of agar (4%) was used which resulted in a semi-solid gel that could stand alone and support implanted leads. Fluoroptic temperature probes (OSENSA, BC, Canada) were attached to the exposed tips of the leads for temperature measurements during RF exposure at 64 MHz and 123 MHz (Figure 6A).

**Figure 5:**
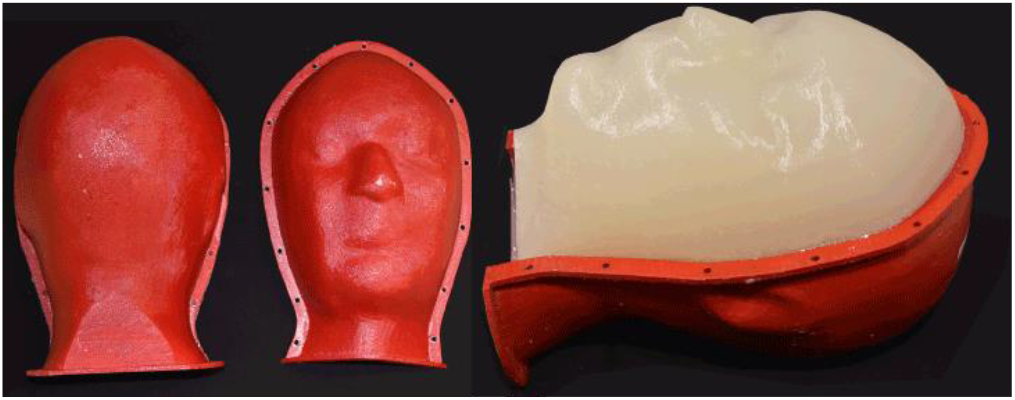
Anthropomorphic head phantom based on MRI of a healthy volunteer

**Figure 6:**
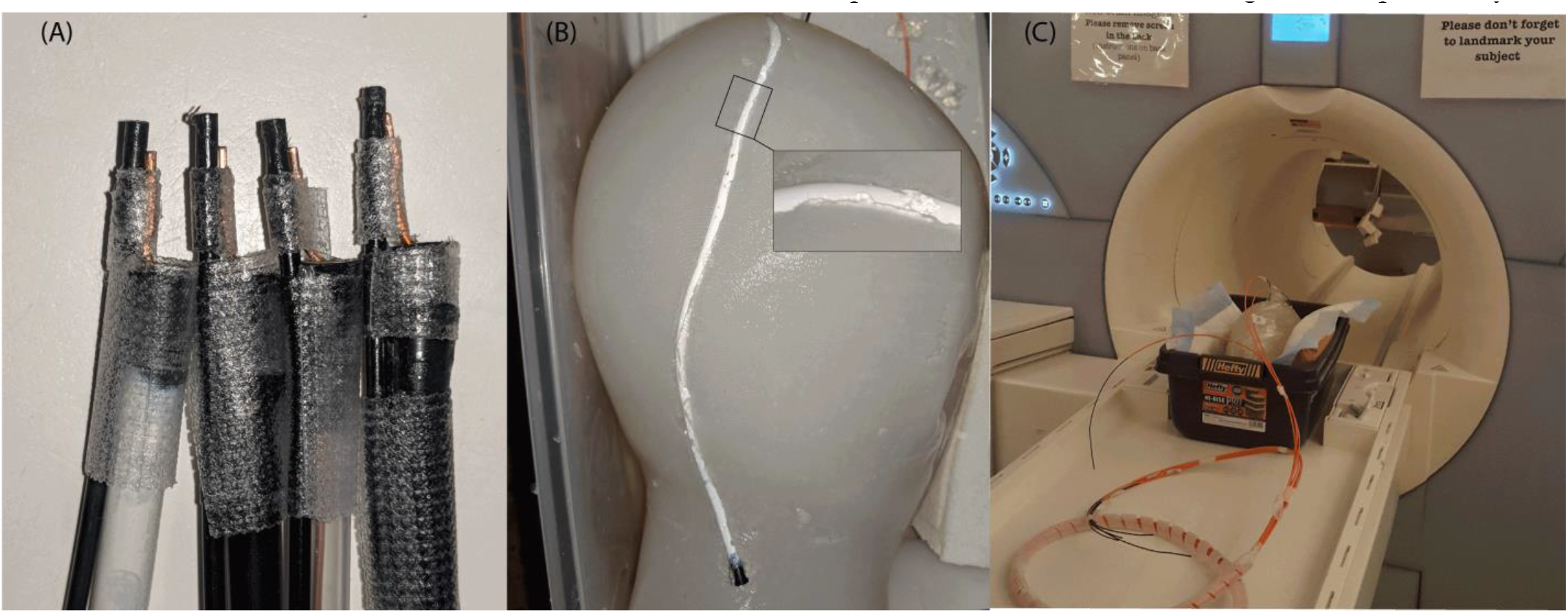
(A) Temperature probes attached to the exposed tips of the leads. Attention was paid to maintain a consistent distance between the tip of the probe and the tip of leads in all four experiments (B) Trajectory of the lead implanted into the gel phantom. Leads were pressed into the gel to form a good galvanic connection with the conductive medium (C) Positioning of the phantom and implants in the MRI system.

It is well known that the trajectory of the implanted lead has a substantial effect on the heating at the tip [32, 33, 43–45]. Therefore, attention was paid to keep the lead trajectories and the phantom positions the same across all experiments. For each experiment, the lead was gently pressed into the gel (~ 1mm deep) to assure a good contact was maintained with the conductive medium (Figure 6B).

Experiments were performed at a 1.5 T Magnetom Avanto system, a 3 T TIM Trio system (Siemens Healthineers, Erlangen, Germany). To have better control over the characteristics of the RF exposure, gradient coils were disabled and a train of 1 ms rectangular RF pulses was transmitted using the scanner body coil. Pulse sequence parameters are summarized in Table I for each experiment.

**TABLE I.** CHARACTERISTICS OF RF PULSES

| Frequency | Flip Angle | Duration |
| --- | --- | --- |
| 64 MHz | 200° | 207 s |
| 123 MHz | 250° | 148 s |

### C. Temperature Measurement Results

Figure 7 gives the measured temperature rise near the tip of each implant after ~2.5 minutes of RF exposure at 1.5 T and 3 T. As predicted, CBLoC lead generated substantially reduced heating near the tip compared to the control lead at both 1.5 T and 3 T, showing a ~20-fold reduction in temperature rise. The results further confirmed our hypothesis that the use of HDC material when the path to the conductive tissue is blocked does not produce the same heat-reducing effect. Specifically, the experiment with non-conductive rubber tubing filled with the HDC material increased the heating at 1.5 T, although some heat-reduction effect was observed at higher fields. Similarly, conductive tubing alone without the HDC material did not produce the same level of heat reduction. It is noteworthy to mention however, that the presence of the conductive tubing alone reduced the heating at 3T to some extent. This effect is most probably attributable to a different mechanism recently observed and termed as “decoy” [46].

**Figure 7:**
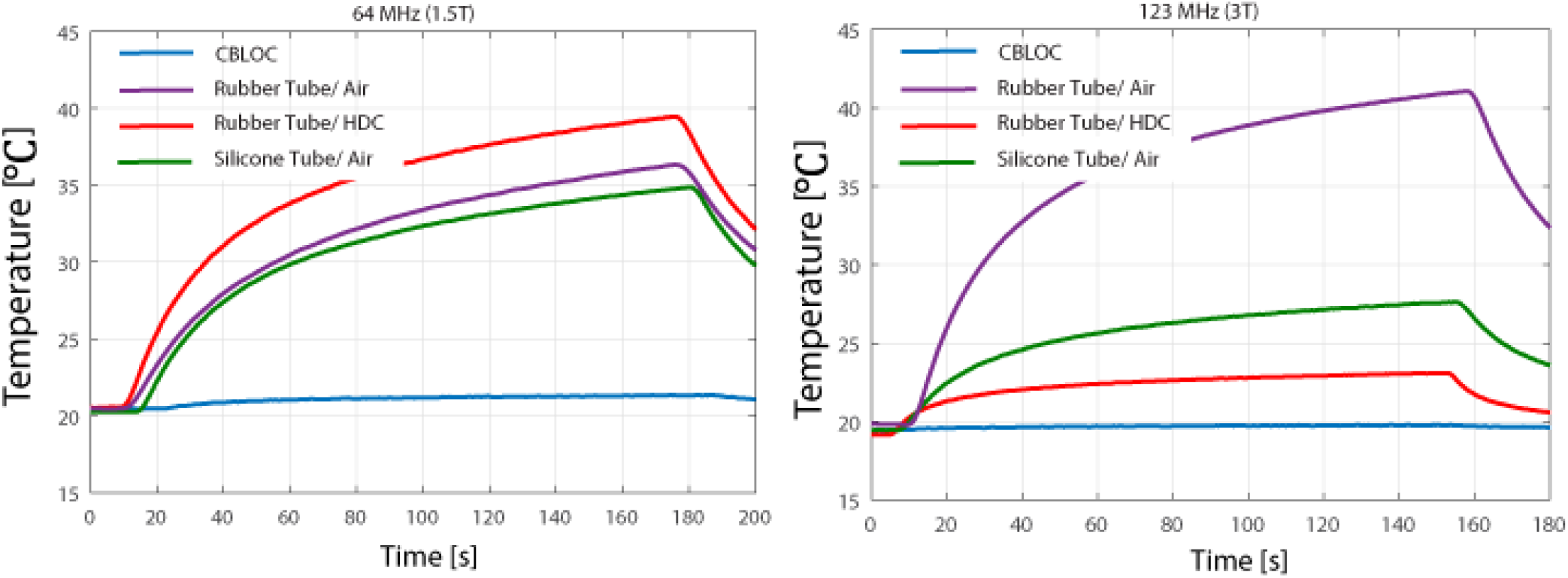
Measured temperature increases near the tip of implanted leads during ~2.5 minutes RF exposure at 64 MHz (1.5T) and 123 MHz (3T).

## IV. SIMULATIONS

A major difficulty in devising a systematic methodology to estimate MRI RF-induced implant heating is that the problem has a large parameter space. In general, the performance of CBLOC leads depends on the interplay between plurality of factors, including frequency and geometry of MRI RF transmitter, thickness and electric properties of HDC coating and insulator, as well as the trajectory of the lead in the phantom. The complexity of RF heating phenomenon and its large parameter space has been emphasized in experimental studies [43, 47]. The application of electromagnetic simulations to better understand the phenomenology of electric and magnetic field interaction with the tissue has proven to be indispensable [32, 48–61]. One exquisite advantage of computational models is that they allow for a controlled and systematic change of design parameters and enable the assessment of multitude of design scenarios in a reasonable time. When using computer models to guide the design however, it is important to verify that the model represents the physical system well enough to be able to reliably predict its behavior for different design parameters. In this section, we present results of numerical simulations calculating the 1g-averaged SAR around the tip the leads that were used in our experiments. Simulations were performed with models of whole-body birdcage transmit coils tuned at 64 MHz (Siemens Avanto system) and 123 MHz (Siemens Tim Trio system). Notably, the goal was not to reproduce the exact amount of heating observed in the experiments, but rather to show that the simulated local SAR is a reliable surrogate to compare the behavior of different leads. We also implemented model of a 7 T head-only coil based on a prototype built in our lab to investigate behavior of CBLOC leads at higher fields.

### A. RF Coils and Lead Models

ANSYS Electronics Desktop (HFSS 16.2, Designer, ANSYS Inc., Canonsburg, PA) was used to implement models of shielded 16-rung high-pass birdcage body coils tuned to 64 MHz (1.5 T proton imaging) and 123 MHz (3T proton imaging). The 7 T head coil was a hybrid birdcage with 16 legs and 128 capacitors distributed around end rings and on the legs. All coils were fed by a quadrature excitation implemented at two ports on one of the end-rings 90° apart in position and phase as illustrated in Figure 8. The details of coils dimensions and values of tuning capacitors are also given in Figure 8.

**Figure 8:**
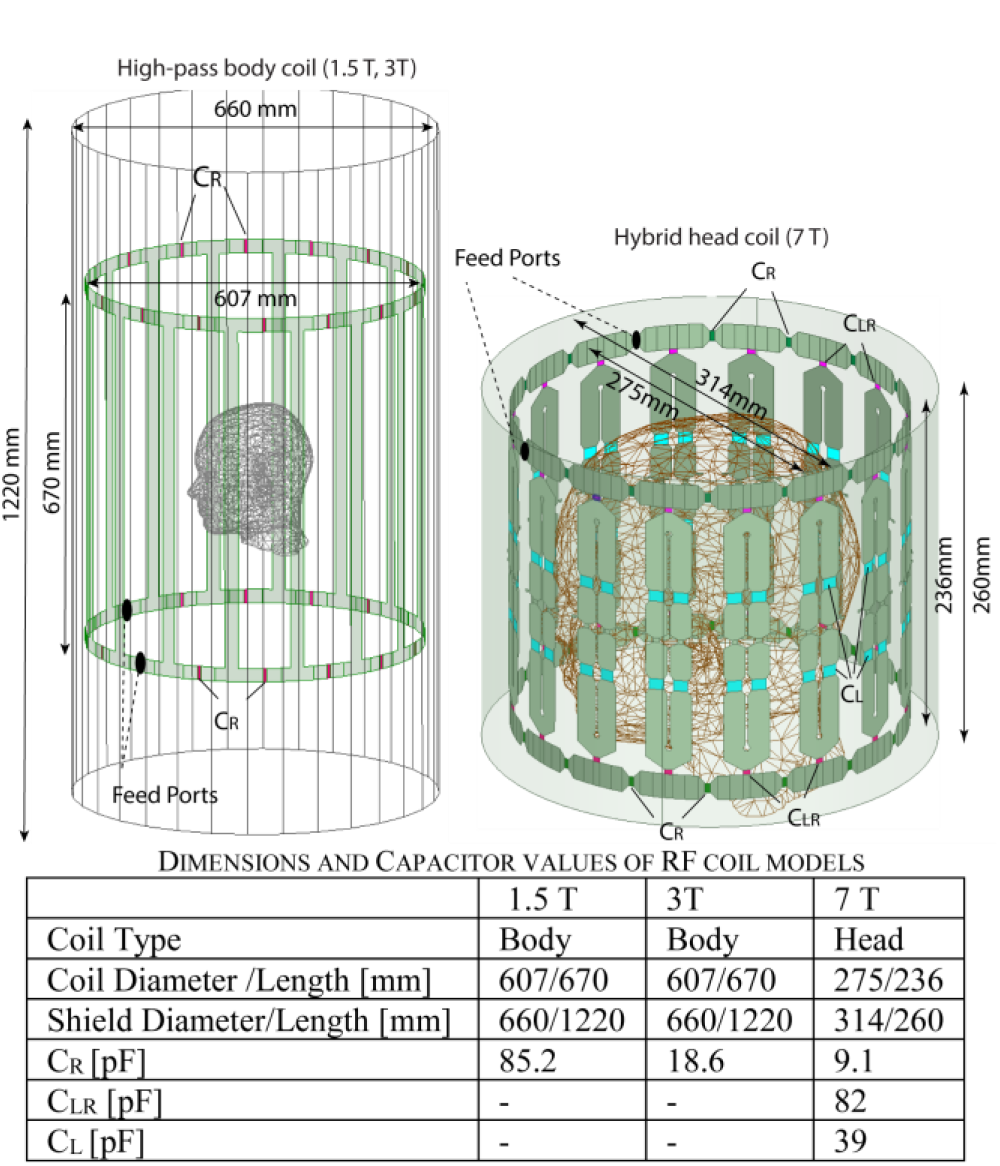
Numerical models of high-pass MRI body coils and a hybrid head coil implemented in ANSYS HFSS

Lead models were constructed and fitted into a homogeneous human head phantom (*ε_r_* = 70, *σ* = 1 *S*/*m*) such that they mimicked the experimental setup as closely as possible. Figure 9 shows the details of the meshed model. For each simulation, the final mesh was visually inspected to ensure that fine features of the leads including the thin layer of varnish on copper wires were appropriately represented. The final model included ~5 million tetrahedral elements and took approximately three hours to run on a Dell PowerEdge R730 system with 16×2GB =512GB of RAM, and 28 cores (2×Intel Xeon CPU with each 14 cores) running 64-bit Windows Server 2012 system.

**Figure 9:**
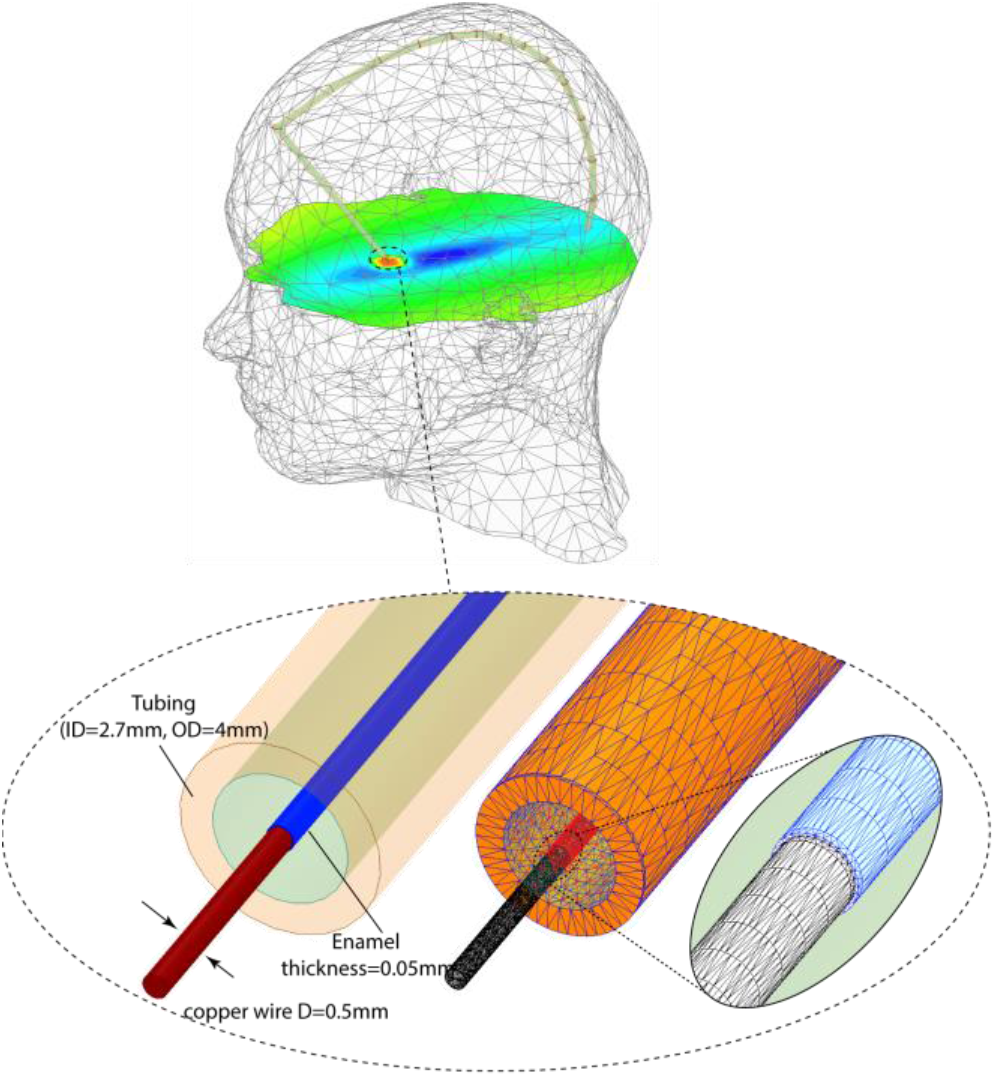
Numerical models of the leads fitted into the human head. The mesh was visually inspected for all simulations to ensure fine features of the leads were properly represented.

### B. Simulation Results

For all simulations, the input power of the coil was adjusted to produce an average B1^+^=2 μT on a transverse plan passing through the center of the head phantom.

The magnitude of the 1g-averaged SAR in a cubic area of 2cm×2cm×2cm surrounding the lead’s tip is given in table III. As it can be observed, the simulated SAR in the tissue around the tip correlated very well with the measured temperature rise at the same location at both 1.5 T and 3 T.

**TABLE II.**
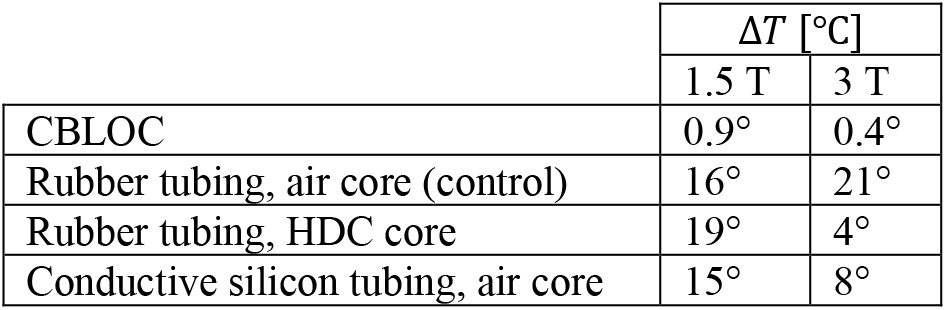
MEASURED TEMPERATURE INCREASE NEAR THE TIP OF DIFFERENT LEADS

At 1.5 T the maximum temperature rise was measured to be 19°C at the tip of the lead with rubber tubing filled with HDC, followed by 16°C at the tip of the lead with empty rubber tubing (control lead), and 15°C at the tip of the lead with empty conductive silicon tubing. The CBLOC lead showed the minimum temperature rise, measured to be just below 1°C. Simulations predicted the same trend in the calculated 1g-averaged SAR, with the maximum of 118 W/kg at the tip of the lead with rubber tubing filled with HDC, 85 W/kg around the tip of the lead with empty rubber tubing, and 82 W/kg at the tip of the lead with empty conductive silicon tubing. The CBLOC lead generated the minimum SAR in the tissue (8 W/kg).

Similarly, at 3 T the calculated SAR perfectly predicted the heating of the leads compared to each other. The maximum temperature rise at 3 T was observed in lead with empty rubber tubing (21°C), followed by the lead with empty conductive silicone tubing (8°C) and the lead with rubber tubing filled with the HDC (4°C). The CBLOC lead generated the minimum heating (< 1 °C). The same trend was observed in the calculate SAR as given in Table III.

**TABLE III.** SIMULATED MAX_SAR1g (W/kg) AT 1.5 T, 3 T AND 7 T.

|  | 1.5 T | 3 T | 7 T |
| --- | --- | --- | --- |
| CBLOC | 8 | 7 | 22 |
| Rubber tubing, air core (control) | 85 | 53 | 336 |
| Rubber tubing, HDC core | 118 | 10 | 48 |
| Conductive silicon tubing, air core | 82 | 20 | 213 |

Simulations at 7 T predicted the same trend as observed in 3T, with the lead with empty rubber tubing producing the largest SAR, followed by the lead with empty conductive silicon tubing and the lead with rubber tubing filled with the HDC. The SAR produced by the CBLOC lead was ~15-fold less than the SAR of the control lead.

## V. DISCUSSION

With the rapid growth of clinical applications of high-field MRI, there has been a significant effort to alleviate the problem of RF heating of implanted conductive leads. This work presents a novel technique to improve MRI RF safety of implanted leads through capacitive bleeding of currents along the length of the lead. The technique termed CBLOC, uses a layer of high-dielectric constant material coating the lead wires and forming distributed capacitance along the length of the lead. This distributed capacitance will dissipate the RF energy along the length of the lead in the form of displacement currents and thus, reduces the heating at the tip.

High permittivity materials have been long used in the electronics industry for manufacturing on-chip capacitors [62–64]. Their application in the context of MRI however, is relatively new. Pioneering work has been done by the Webb group advancing the use of HDC pads to increase the signal to noise ratio during high-field MRI [42, 65–67]. Recently, the use of external HDC pads in regions not directly adjacent to the implant cite has been shown to reduce the SAR at the tip of medical implants [68]. In our experiments, we observed a substantial reduction of the heating at the tip of CBLOC implants by up to 20-fold during 1.5 T and up to 40-fold at 3 T MRI compared to the control case. Simulation studies calculating 1g-averaged SAR around the tip of implanted leads predicted the trend of observed heating at both 1.5 T and 3T. Simulations with head-only coils at 7T predicted a similar performance. From the practical point of view, the real-life implementation of such leads should be possible using conventional techniques for atomic layer deposition of high-dielectric oxides on conductive wires [69, 70]. In such case, the solid HDC layer can act as the insulation for the lead wires, eliminating the need to use tubing.

The current study has a few limitations. First, we examined the performance of CBLOC leads inside a homogeneous head phantom, thus the effect of tissue heterogeneity was not studied. Future studies should evaluate the effect of CBLOC in realistic multi-compartment body models. Furthermore, it is well established that implant’s trajectory has a non-negligible effect on the heating, as it directly affects the coupling of RF fields with conductive wires. Thus, more studies are required to analyze sensitivity of the results to factors such as composition of surrounding tissue and trajectory of the implant. Finally, the effect of adding the HDC coating on the mechanical stability and flexibility of the leads needs to be evaluated.

## VI. CONCLUSION

A new promising approach to reduce the heating of implanted conductive leads during MRI scans has been presented. Both experimental results and numerical simulations are presented to demonstrate the substantial heat reduction performance of the new lead. The technique is based on capacitive shunting of RF energy along the length of the lead through the high-dielectric constant material. The proposed approach may lead to the design of a new generation of implantable leads for MRI.

## VII. Acknowledgment

This work was supported by the NIH grant R03EB024705.

Disclaimer: The mention of commercial products, their sources, or their use in connection with material reported herein is not to be construed as either an actual or implied endorsement of such products by the Department of Health and Human Services

---

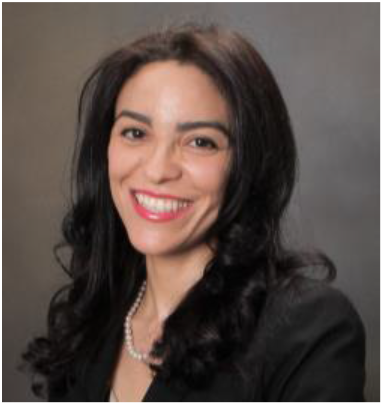
Laleh Golestanirad received her PhD in electrical engineering with a focus on computational electromagnetics from Ecole Polytechnique Federal de Lausanne (EPFL), Switzerland, in 2011. She pursued further training in magnetic resonance imaging (MRI) hardware instrumentation and safety assessment as a postdoctoral fellow at Department of Medical Biophysics, University of Toronto and A. A. Martinos Center for Biomedical Imaging, Harvard Medical School. In 2016 she received NIH Career Award K99/R00 to develop MRI methodologies for safe imaging of patients with deep brain stimulation implants. She then joined the Department of Biomedical Engineering at Northwestern University as an Assistant Professor in 2018 where she works on developing MRI technologies for imaging of neural implants and application of computational modeling in brain stimulation techniques.

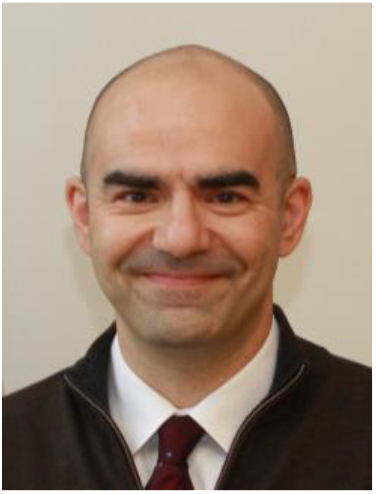
Leonardo M. Angelone (M’04) is a Research Biomedical Engineer at the Office of Science and Engineering Laboratories, Center of Devices and Radiological Health, U.S. FDA. Dr. Angelone leads a Research program that investigates assessment of energy deposition and heating induced in the human body by medical devices using electromagnetic energy. The results of the projects have been presented in over 100 peer-reviewed journal articles and conference proceedings, and publicly available software. Dr. Angelone completed a Laurea in Electronic Engineering (University “La Sapienza”, Rome, Italy), a Ph.D. in Biomedical Engineering (Tufts University, Medford, MA), and a Research Fellowship at the A. Martinos Center for Biomedical Imaging, Department of Radiology of the Massachusetts General Hospital, Harvard Medical School. Prior to joining the FDA, Dr. Angelone has been a consultant with the Research and Development Department in the Surgical Products Division of Hologic Inc.

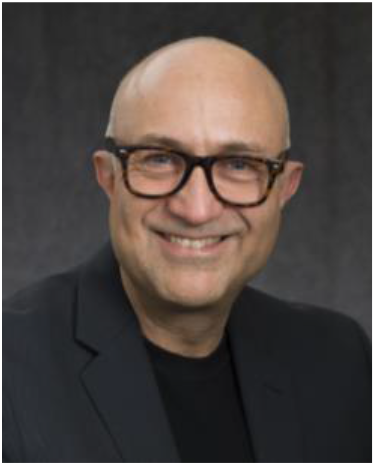
Giorgio Bonmassar (S’89, M’90, SM’10) was born in Milan, Italy, on May 13, 1962. He received the Laurea degree in EE from the University of Rome “La Sapienza,” Rome, Italy, in 1989, and the Ph.D. degree in biomedical engineering from Boston University, Boston, MA, in 1997. From 1989 to 1991, he was a Resea rch and Development Systems Engineer with Ericsson, Rome, Italy, a Research Fellow (1992–1997) and a Post-Doctoral Fellow (1997) with Boston University, Boston, MA, a Research Fellow (1998–2000) with Massachusetts General Hospital, Boston, MA and an Instructor (2000–2005) with Massachusetts General Hospital, Boston, MA. In 2005, he became an Assistant Professor and since 2017 he is an Associate Professor in Radiology with the AA. Martinos Center, Massachusetts General Hospital, Harvard Medical School, Charlestown, MA. He has authored or coauthored over 100 international journal papers and conference presentations on biomedical engineering. Dr. Bonmassar is a member of the International Society for Magnetic Resonance Imaging and the Alfa Eta Mu Beta Biomedical Engineering Research Society. He was the recipient of a 1999 North American Treaty Organization (NATO) Advanced Research Studies Award and a 2000 Whitaker Foundation Biomedical Engineering Grant for Young Investigators. Since 2003, he is and has been the principal investigator (PI) on a many National Institutes of Health (NIH) and Department of Defense (DoD) grants.

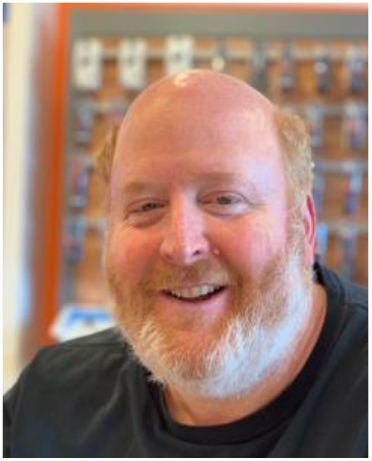
Lawrence L. Wald, Ph.D., is currently a Professor of Radiology at Harvard Medical School and Affiliated Faculty of the Harvard-MIT Division Health Sciences Technology. He received a BA in Physics at Rice University, and a Ph.D. in Physics from the University of California at Berkeley in 1992 under the direction of Prof. E.L. Hahn with a thesis related to optical detection of NMR. He obtained further (postdoctoral) training in Physics at Berkeley and then in Radiology and MRI at the University of California at San Francisco (UCSF). He began his academic career as an Instructor at the Harvard Medical School at McLean Hospital and since 1998 has been at the Massachusetts General Hospital Dept. of Radiology in the NMR Center (now the A.A. Martinos Center for Biomedical Imaging). His recent work has explored the benefits and challenges of highly parallel MRI and its application to highly accelerated image encoding and parallel excitation, ultra-high field MRI (7 Tesla) methodology for brain imaging, improved method for studying the Human Connectome and portable MRI technology.

## References

[1] A. J. Greenspon et al., “Trends in permanent pacemaker implantation in the United States from 1993 to 2009: increasing complexity of patients and procedures,” Journal of the American College of Cardiology, vol. 60, no. 16, pp. 1540–1545, 2012.

[2] T. Sommer et al., “Strategy for Safe Performance of Extrathoracic Magnetic Resonance Imaging at 1.5 Tesla in the Presence of Cardiac Pacemakers in Non-Pacemaker-Dependent Patients A Prospective Study With 115 Examinations,” Circulation, vol. 114, no. 12, pp. 1285–1292, 2006.

[3] “Neurostimulation Devices Market Size, Share, Growth Report, 2023: Accessed Online January 2018 https://www.gminsights.com/industrv-analYsis/neurostimulation-devices-market-report,” 2016.

[4] R. Kalin and M. S. Stanton, “Current clinical issues for MRI scanning of pacemaker and defibrillator patients,” Pacing and clinical electrophysiology, vol. 28, no. 4, pp. 326–328, 2005.

[5] C. P. Naehle et al., “Evaluation of cumulative effects of MR imaging on pacemaker systems at 1.5 Tesla,” Pacing and clinical electrophysiology, vol. 32, no. 12, pp. 1526–1535, 2009.

[6] D. Dormont, D. Seidenwurm, D. Galanaud, P. Cornu, J. Yelnik, and E. Bardinet, “Neuroimaging and deep brain stimulation,” American Journal of Neuroradiology, vol. 31, no. 1, pp. 15–23, 2010.

[7] A. R. Rezai et al., “Neurostimulation systems for deep brain stimulation: In vitro evaluation of magnetic resonance imaging-related heating at 1.5 tesla,” Journal of Magnetic Resonance Imaging, vol. 15, no. 3, pp. 241–250, 2002.

[8] W. R. Nitz, A. Oppelt, W. Renz, C. Manke, M. Lenhart, and J. Link, “On the heating of linear conductive structures as guide wires and catheters in interventional MRI,” Journal of Magnetic Resonance Imaging, vol. 13, no. 1, pp. 105–114, 2001.

[9] J. M. Henderson, J. Tkach, M. Phillips, K. Baker, F. G. Shellock, and A. R. Rezai, “Permanent neurological deficit related to magnetic resonance imaging in a patient with implanted deep brain stimulation electrodes for Parkinson’s disease: case report,” Neurosurgery, vol. 57, no. 5, p. E1063, 2005.

[10] J. Spiegel et al., “Transient dystonia following magnetic resonance imagingin a patient with deep brain stimulation electrodes for the treatment of Parkinson disease: Case report,” Journal of neurosurgery, vol. 99, no. 4, pp. 772–774, 2003.

[11] MRI Guidelines for Medtronic Deep Brain Stimulation Systems (http://manuals.medtronic.com/wcm/groms/mdtcomsg/@emanuals/@era/@neuro/documents/documents/contrib228155.ydf), 2015 Date of access: 12/09/2015.

[12] Medtronic, “MRI guidelines for Medtronic neurostimulation systems for chronic pain (http://www.mrisurescan.com/wcm/groups/mdtcomsg/@emanuals/@era/@neuro/documents/documents/contrib161416.pdf),” 2017.

[13] Y. Eryaman, E. A. Turk, C. Oto, O. Algin, and E. Atalar, “Reduction of the radiofrequency heating of metallic devices using a dual-drive birdcage coil,” Magnetic Resonance in Medicine, 2012.

[14] Y. Eryaman, B. Akin, and E. Atalar, “Reduction of implant RF heating through modification of transmit coil electric field,” Magnetic Resonance in Medicine, vol. 65, no. 5, pp. 1305–1313, 2011.

[15] L. Golestanirad, B. Keil, L. M. Angelone, G. Bonmassar, A. Mareyam, and L. L. Wald, “Feasibility of using linearly polarized rotating birdcage transmitters and close-fitting receive arrays in MRI to reduce SAR in the vicinity of deep brain simulation implants,” Magnetic resonance in medicine, vol. 77, no. 4, pp. 1701–1712, 2017.

[16] L. Golestanirad et al., “Construction and modeling of a reconfigurable MRI coil for lowering SAR in patients with deep brain stimulation implants,” Neuroimage, vol. 147, pp. 577–588, 2017.

[17] C. McElcheran, B. Yang, K. J. Anderson, L. Golenstani-Rad, and S. J. Graham, “Investigation of Parallel Radiofrequency Transmission for the Reduction of Heating in Long Conductive Leads in 3 Tesla Magnetic Resonance Imaging,” PLoS One, vol. 10, no. 8, p. e0134379, 2015.

[18] C. E. McElcheran, B. Yang, K. J. Anderson, L. Golestanirad, and S. J. Graham, “Parallel radiofrequency transmission at 3 tesla to improve safety in bilateral implanted wires in a heterogeneous model,” Magnetic resonance in medicine, vol. 78, no. 6, pp. 2406–2415, 2017.

[19] E. Atalar, J. Allen, P. Bottomley, W. Eldelstein, and P. V. Karmarkar, “MRI-safe high impedance lead systems,” US8055351B2, 2011.

[20] S. Denker, A. J. Beutler, and C. Bulkes, “MRI compatible implanted electronic medical device,” US8255054B2, 2012.

[21] J. M. Olsen, G. A. Hrdlicka, C. D. Wahlstrand, and T. B. Hoegh, “Lead electrode for use in an MRI-safe implantable medical device,” US7853332B2, 2010.

[22] R. A. Stevenson et al., “Implantable lead bandstop filter employing an inductive coil with parasitic capacitance to enhance MRI compatibility of active medical devices,” US8145324B1, 2012.

[23] A. Vase and D. N. Sethna, “Implantable medical lead configured for improved mri safety,” US20090270956A1, 2009.

[24] E. Villaseca and G. Dublin, “Electromagnetic trap for a lead,” US20030144720A1, 2003.

[25] C. D. Wahlstrand, T. B. Hoegh, G. A. Hrdlicka, T. E. Cross Jr, and J. M. Olsen, “Lead electrode for use in an MRI-safe implantable medical device,” US7174219B2, 2007.

[26] V. Zeijlemaker, “Method and apparatus for shielding coating for MRI resistant electrode systems,” US20030144718A1, 2003.

[27] P. Serano, L. M. Angelone, H. Katnani, E. Eskandar, and G. Bonmassar, “A novel brain stimulation technology provides compatibility with MRI,” Scientific reports, vol. 5, p. 9805, 2015.

[28] P. A. Bottomley, P. V. Karmarkar, J. M. Allen, and W. A. Edelstein, “MRI and RF compatible leads and related methods of operating and fabricating leads,” US20080243218A1, 2016.

[29] P. A. Bottomley, A. Kumar, W. A. Edelstein, J. M. Allen, and P. V. Karmarkar, “Designing passive MRI-safe implantable conducting leads with electrodes,” Medical physics, vol. 37, no. 7, pp. 3828–3843, 2010.

[30] S. McCabe and J. Scott, “A novel implant electrode design safe in the rf field of mri scanners,” IEEE Transactions on Microwave Theory and Techniques, vol. 65, no. 9, pp. 3541–3547, 2017.

[31] R. Das and H. Yoo, “RF Heating Study of a New Medical Implant Lead for 1.5 T, 3 T, and 7 T MRI Systems,” IEEE Transactions on Electromagnetic Compatibility, vol. 59, no. 2, pp. 360–366, 2017.

[32] L. Golestanirad, L. M. Angelone, M. I. Iacono, H. Katnani, L. L. Wald, and G. Bonmassar, “Local SAR near deep brain stimulation (DBS) electrodes at 64 MHz and 127 MHz: A simulation study of the effect of extracranial loops” Magnetic Resonance in Medicine vol. 88, no. 4, pp. 1558–1565, 2016.

[33] K. B. Baker, J. Tkach, J. D. Hall, J. A. Nyenhuis, F. G. Shellock, and A. R. Rezai, “Reduction of magnetic resonance imaging-related heating in deep brain stimulation leads using a lead management device,” Neurosurgery, vol. 57, no. 4, pp. 392–397, 2005.

[34] S. M. Park, R. Kamondetdacha, and J. A. Nyenhuis, “Calculation of MRI-induced heating of an implanted medical lead wire with an electric field transfer function,” Journal of Magnetic Resonance Imaging, vol. 26, no. 5, pp. 1278–1285, 2007.

[35] C. J. Yeung, R. C. Susil, and E. Atalar, “RF heating due to conductive wires during MRI depends on the phase distribution of the transmit field,” Magnetic Resonance in Medicine, vol. 48, no. 6, pp. 1096–1098, 2002.

[36] R. Buchli, P. Boesiger, and D. Meier, “Heating effects of metallic implants by MRI examinations,” Magnetic resonance in medicine, vol. 7, no. 3, pp. 255–261, 1988.

[37] C.-K. Chou, J. A. McDougall, and K. W. Chan, “RF heating of implanted spinal fusion stimulator during magnetic resonance imaging,” IEEE transactions on biomedical engineering, vol. 44, no. 5, pp. 367–373, 1997.

[38] C. Bulkes and S. Denker, “Mri compatible implanted electronic medical device and lead,” 2007.

[39] E. Atalar, J. Allen, P. Bottomley, W. Eldelstein, and P. V. Karmarkar, “MRI-safe high impedance lead systems,” ed: Google Patents, 2006.

[40] R. A. Stevenson et al., “Implantable lead bandstop filter employing an inductive coil with parasitic capacitance to enhance MRI compatibility of active medical devices,” ed: Google Patents, 2012.

[41] W. K. Chen, The electrical engineering handbook. Academic press, 2004.

[42] W. Teeuwisse, W. Brink, K. Haines, and A. Webb, “Simulations of high permittivity materials for 7 T neuroimaging and evaluation of a new barium titanate-based dielectric,” Magnetic resonance in medicine, vol. 67, no. 4, pp. 912–918, 2012.

[43] E. Mattei et al., “Complexity of MRI induced heating on metallic leads: experimental measurements of 374 configurations,” Biomedical engineering online, vol. 7, no. 1, p. 11, 2008.

[44] P. Nordbeck et al., “Spatial distribution of RF-induced E-fields and implant heating in MRI,” Magnetic resonance in medicine, vol. 60, no. 2, pp. 312–319, 2008.

[45] P. Nordbeck et al., “Measuring RF-induced currents inside implants: Impact of device configuration on MRI safety of cardiac pacemaker leads,” Magnetic resonance in medicine, vol. 61, no. 3, pp. 570–578, 2009.

[46] S. McCabe and J. Scott, “A Novel Implant Electrode Design Safe in the RF Field of MRI Scanners,” IEEE Transactions on Microwave Theory and Techniques, 2017.

[47] G. Calcagnini et al., “In vitro investigation of pacemaker lead heating induced by magnetic resonance imaging: role of implant geometry,” Journal of Magnetic Resonance Imaging, vol. 28, no. 4, pp. 879–886, 2008.

[48] E. Cabot et al., “Evaluation of the RF heating of a generic deep brain stimulator exposed in 1.5 T magnetic resonance scanners,” Bioelectromagnetics, vol. 34, no. 2, pp. 104–113, 2013.

[49] B. L. Wilkoff et al., “Safe magnetic resonance imaging scanning of patients with cardiac rhythm devices: a role for computer modeling,” Heart Rhythm, vol. 10, no. 12, pp. 1815–1821, 2013.

[50] C. M. Collins et al., “Temperature and SAR calculations for a human head within volume and surface coils at 64 and 300 MHz,” Journal of Magnetic Resonance Imaging, vol. 19, no. 5, pp. 650–656, 2004.

[51] L. Golestani-Rad, B. Elahi, and J. Rashed-Mohassel, “Investigating the effects of external fields polarization on the coupling of pure magnetic waves in the human body in very low frequencies,” Biomagnetic research and technology, vol. 5, no. 1, p. 3, 2007.

[52] L. Golestanirad et al., “Combined use of transcranial magnetic stimulation and metal electrode implants: a theoretical assessment of safety considerations,” Physics in medicine and biology, vol. 57, no. 23, p. 7813, 2012.

[53] L. Golestanirad, B. Elahi, A. Molina Arribere, J. R. Mosig, C. Pollo, and S. J. Graham, “Analysis of fractal electrodes for efficient neural stimulation,” Frontiers in neuroengineering, vol. 6, p. 3, 2013.

[54] L. Golestanirad, A. P. Izquierdo, S. J. Graham, J. R. Mosig, and C. Pollo, “Effect of realistic modeling of deep brain stimulation on the prediction of volume of activated tissue,” Progress In Electromagnetics Research, vol. 126, pp. 1–16, 2012.

[55] L. Golestanirad, M. Mattes, J. R. Mosig, and C. Pollo, “Effect of model accuracy on the result of computed current densities in the simulation of transcranial magnetic stimulation,” IEEE Transactions on Magnetics, vol. 46, no. 12, pp. 4046–4051, 2010.

[56] L. Golestanirad et al., “RF-induced heating in tissue near bilateral DBS implants during MRI at 1.5 T and 3T: The role of surgical lead management,” Neuroimage, vol. 184, pp. 566–576, 2019.

[57] P. Wei, B. Yang, C. McElcheran, L. Golestanirad, and S. Graham, “Reducing Radiofrequency-induced Heating in Realistic Deep Brain Stimulation Lead Trajectories using Parallel Transmission,” in Proc. Intl. Soc. Mag. Reson. Med.26, 2018.

[58] L. Navarro de Lara et al., “Combined EM simulations and measurements of birdcage coil B1+ for designing a 3T multichannel TMS /MRI head coil array,” in Proc. Intl. Soc. Mag. Reson. Med. 26, 2018.

[59] L. Golestanirad et al., “Solenoidal micromagnetic stimulation enables activation of axons with specific orientation,” Frontiers in Physiology, vol. 9, no. 724, 2018.

[60] Giorgio Bonmassar, Laleh Golestanirad, and J. Deng, “Enhancing Coil Design for Micromagnetic Brain Stimulation,” MRS Advances vol. 3, no. 9, pp. 1635–1640, 2018.

[61] L. Golestanirad et al., “Variation of RF heating around deep brain stimulation leads during 3.0 T MRI in fourteen patient-derived realistic lead models: The role of extracranial lead management,” in Proc. Intl. Soc. Mag. Reson. Med. 25, 2017.

[62] Y. Rao, S. Ogitani, P. Kohl, and C. Wong, “Novel polymer–ceramic nanocomposite based on high dielectric constant epoxy formula for embedded capacitor application,” Journal of Applied Polymer Science, vol. 83, no. 5, pp. 1084–1090, 2002.

[63] G. Sandhu and P. Fazan, “High dielectric constant capacitor and method of manufacture,” ed: Google Patents, 1994.

[64] Y. Rao and C. Wong, “Material characterization of a high-dielectric-constant polymer–ceramic composite for embedded capacitor for RF applications,” Journal of Applied Polymer Science, vol. 92, no. 4, pp. 2228–2231, 2004.

[65] W. M. Brink, A. M. van der Jagt, M. J. Versluis, B. M. Verbist, and A. G. Webb, “High permittivity dielectric pads improve high spatial resolution magnetic resonance imaging of the inner ear at 7 T,” Investigative radiology, vol. 49, no. 5, pp. 271–277, 2014.

[66] W. M. Brink and A. G. Webb, “High permittivity pads reduce specific absorption rate, improve B1 homogeneity, and increase contrast-to-noise ratio for functional cardiac MRI at 3 T,” Magnetic resonance in medicine, vol. 71, no. 4, pp. 1632–1640, 2014.

[67] B. S. Park, B. McCright, L. M. Angelone, A. Razjouyan, and S. S. Rajan, “Improvement of Electromagnetic Field Distributions Using High Dielectric Constant (HDC) Materials for CTL-Spine MRI: Numerical Simulations and Experiments,” IEEE Transactions on Electromagnetic Compatibility, vol. 59, no. 5, pp. 1382–1389, 2017.

[68] Z. Yu, X. Xin, and C. M. Collins, “Potential for high-permittivity materials to reduce local SAR at a pacemaker lead tip during MRI of the head with a body transmit coil at 3 T,” Magnetic resonance in medicine, vol. 78, no. 1, pp. 383–386, 2017.

[69] S. K. Kim, W.-D. Kim, K.-M. Kim, C. S. Hwang, and J. Jeong, “High dielectric constant TiO 2 thin films on a Ru electrode grown at 250 C by atomic-layer deposition,” Applied Physics Letters, vol. 85, no. 18, pp. 4112–4114, 2004.

[70] Y. Ono, W.-W. Zhuang, and R. Solanki, “Methods of using atomic layer deposition to deposit a high dielectric constant material on a substrate,” ed: Google Patents, 2002.

